# Retention of adults from fishing communities in an HIV vaccine preparedness study in Masaka, Uganda

**DOI:** 10.1101/365148

**Authors:** Ubaldo Bahemuka, Andrew Abaasa, Eugene Ruzagira, Christina Lindan, Matt A Price, Anatoli Kamali, Pat Fast

## Abstract

**Introduction:** People living in fishing communities around Lake Victoria may be suitable for enrolment in HIV prevention trials because of high HIV incidence. We assessed the ability to recruit and retain individuals from fishing communities into an HIV vaccine preparedness cohort study in Masaka, Uganda.

**Methods:** HIV high risk, sero-negative adults (18-49 years) were identified from four fishing villages bordering Lake Victoria through door-to-door HIV counselling and testing (HCT). Interested persons were referred for: screening, enrolment, and quarterly follow-up visits at a study clinic located approximately 40 kilometres away. Repeat HCT, HIV risk assessment, and evaluation and treatment for sexually transmitted infections were provided. Rates of and factors associated with study dropout were assessed using Poisson regression models.

**Results:** A total of 940 participants were screened between January 2012 and February 2015, of whom 654 were considered for the analysis. Over a two-year follow-up period, 197 (30.1%) participants dropped out of the study over 778.9 person-years, a dropout rate of 25.3 / 100 person-years. Dropout was associated with being female (aRR =1.56, 95% confidence interval [CI] 1.12-2.18), age, being 18-24 years (aRR=1.64; 95% CI 1.03-2.60), 25-34 years (aRR=1.63; 95% CI 1.04-2.55); having no education (aRR=2.02; 95% CI: 1.23-3.31); living in the community for less than one year (aRR=2.22; 95% CI: 1.46-3.38) or 1-5 years (aRR=1.68; 95% CI: 1.16-2.45) and occupation.

**Conclusions:** It is possible to recruit and retain individuals from fishing communities, however, intensified participant tracing may be necessary in a vaccine trial to keep in follow up female, young, less educated, those in mobile occupations and new residents.

## Introduction

New HIV infections continue to occur in sub-Saharan Africa, despite behavioural and biomedical prevention efforts(1). Therefore, development of an effective HIV vaccine will be essential to reducing incidence in this region(2). Efficacy testing of HIV vaccine candidates is more efficient among populations with favourable retention and high HIV incidence rates(3). Fishing communities in Uganda have a high burden of HIV with HIV prevalence and incidence ranging from 20-40 percent (4–6) and 3-9 cases per 100 person-years of observation (PYO) respectively (7–9). This high level of risk suggests that persons living in these communities may be an ideal population for HIV vaccine efficacy trials. However, fishing communities are typically located in remote areas and often have little if any health care infrastructure, factors that may present challenges to the rigorous implementation requirements of HIV vaccine preparedness and efficacy studies(10).

A number of studies have evaluated the suitability of fishing communities around Lake Victoria for participation in future HIV vaccine efficacy trials in terms of participant retention. In three recent studies, (9, 11, 12) the 12-month retention was reported to be 83%, 77% and 85% respectively. In these studies, recruitment and follow up assessments were conducted at clinics established in each of the participating fishing community. During the conduct of a Simulated HIV vaccine trial(SiVET) in the cohort of our study, Abaasa et al (13)reported a 73%% retention in the observational arm of the study.

In some cases, it may be advantageous to conduct an efficacy trial in a central location, recruiting volunteers from surrounding areas. The clinical research centre in Masaka town situated is inland from the neighbouring fishing communities. Little is known about how retention might be affected if participants from fishing communities had to travel to a clinic outside of their communities to attend study visits. In order to answer this question, we investigated retention and dropout rates among fisher folk enrolled in a HIV vaccine preparedness cohort at a research clinic located approximately 40 km away from their communities.

## Methods

### Setting and participant identification

The International AIDS Vaccine Initiative (IAVI) in collaboration with the Medical Research Council/Uganda Virus Research Institute Research (MRC/UVRI) has supported a clinical research centre in rural South Western Uganda with the aim of conducting future HIV vaccine efficacy trials. The research Centre is located in Masaka town, approximately 30 to 40 kilometers (km) away from the shores of Lake Victoria. The clinic is equipped to conduct HIV vaccine trials, Good Clinical Laboratory Practice (GCLP) accredited laboratory, a vaccine pharmacy, and Good Clinical Practice (GCP) trained staff (14). The HIV vaccine preparedness study described here was conducted among individuals identified from four mainland fishing communities on the shore of Lake Victoria located in Masaka and Kalungu districts, in southwest Uganda. These communities were selected because they had the largest populations (≥1000 adults) among fishing communities in these two districts.

Between January 2012 and January 2015 a field study team consisting of both male and female counselors visited houses, docked boats, fishing stalls and other venues at the four landing-sites, to offer free rapid HIV testing and counseling on-site to adults. Persons aged 18-49 years old who were identified as HIV uninfected were asked if they would be willing to enroll into a longitudinal study that would require repeated travel to the research clinic. Those who had a positive rapid test result underwent confirmatory testing by having venous blood samples re-tested at the MRC/UVRI laboratory in Entebbe. The participants with confirmed HIV positive results were invited by the field team on phone or physically to receive their results at the clinic approximately one to two weeks after the blood draw. HIV positive participants were linked to care at a treatment center of their choice in accordance with the clinics HIV referral plan.

### Screening and enrolment

Interested individuals who travelled to the study clinic were given detailed information about study procedures in a group setting. Individuals also had one-on-one sessions with a study nurse-counselor who provided additional study information, answered questions, and obtained written informed consent. After obtaining consent participants were assessed for eligibility based on the following criteria: being 18-49 years, HIV uninfected, and sexually active defined as having had sex at least once in the last the three months. Participants also needed to be considered at high risk of HIV acquisition by reporting at least one of the following: self-reported history of a sexually transmitted infection (STI) in the last three months, or the presence of an STI based on medical history/physical exam; condomless vaginal or anal intercourse with a new or with more than one sexual partner in the past 3 months; staying away from home for at least two nights in the past three months; drinking alcohol at least once a week or using illicit drugs (marijuana, Khat, or any other stimulants) in the past month. Participants who fulfilled the above criteria were enrolled into the study. They then completed interviewer-administered questionnaires in either the local language (Luganda) or English, depending on their preference. All enrolled participants had a baseline demographic interview and HIV risk assessment by the study nurse counsellor. The study physician, performed a medical history and physical exam to assess for STI and circumcision status for males. Participants provided physical addresses and phone numbers in order to facilitate future contacts and follow-up.

### Follow-up visits

Participants returned to the clinic every three months for repeat HIV counselling and testing, provision of updated locator information and interim medical history. Everyone underwent a symptom-directed physical exam and a clinical evaluation for STIs. Every six months’ participants provided information on HIV risk behaviour, and were evaluated annually if they continued to fulfil the requirements for participation in the study. At month 12, volunteers were re-assessed for eligibility based on the above risk of HIV acquisition criteria.

Participants who were determined to no longer be eligible were withdrawn from the study. If a participant missed a scheduled appointment, study staff would attempt to contact them via phone or physical tracing by visiting their home. A group of 10 study participants who were considered peer leaders assisted with participant tracing. Participants who missed two consecutive visits were considered to have dropped-out at the time of their last study visit. To bolster study retention, continuous interaction was kept with community advisory board and study staff conducted a series of community activities (research awareness meetings and football matches among others) as a way of improving research awareness. Participants received reimbursement for transportation and time (5000 Uganda shillings, approximately 1.4 USD) at the end of each visit.

### Laboratory testing

Rapid HIV antibody testing was performed on venous blood samples using a single rapid test, Alere Determine (Alere Medical Company Ltd. Chiba, Japan). Specimens that were positive by rapid test were tested with two enzyme linked immunosorbent assay (ELISA) tests in parallel (Murex Biotech Limited, Dartford, United Kingdom, and Vironostika, BioMérieux boxtel, The Netherlands). Discrepant ELISA test results were resolved by testing with either Statpak (Chembio Diagnostic Systems Inc., USA) or Western Blot (Cambridge Biotech, USA). The results of HIV negative rapid tests were given to participants immediately; while participants with confirmed HIV positive results were invited by the field team to receive their results approximately one to two weeks after the blood draw and referred as described earlier. Syphilis testing was performed using the rapid plasma reagin (RPR) test (Microvue, Becton Dickson, Maryland, USA). Samples with an RPR titre of >=1:8 were also tested using the Treponema Pallidum Hemaglutination Assay (TPHA) (Biotech Laboratories, UK). Active syphilis infection was defined as having both a positive RPR titre of >=1:8 and a positive TPHA result.

### Statistical analysis

Data were analysed in Stata 14.0 (Stata Corp, College Station, TX, USA). Participant baseline characteristics were summarised using proportions and means, and stratified by gender. Proportions and means were used to compare baseline characteristics of participants who dropped out of the study to those that completed their scheduled follow up visits. Dropout was defined as all-cause (including withdrawal by self or investigator, death, unknown loss-to-follow-up and refusal to continue) over a maximum of 24 months of follow-up. Participants that were ineligible at the annual reassessment of eligibility to continue were not considered as dropouts and contributed follow up time up to 12 months. HIV sero-positive participants were not considered to have dropped out, but their follow up was censored at the estimated date of HIV infection. Time of HIV infection was estimated as the midpoint between the date of the last negative and the first positive HIV test result. Seventy-two of the enrolled participants that were enrolled but did not return for any follow up visit had their person years of observation (PYO) corrected to one month and the same added to all participants’ PYO. The dropout rate was estimated as the number of participants who dropped out divided by the total person years of observation (PYO). The PYO were calculated as the sum of the time from enrolment (baseline) to the date of the last clinic visit or date of censoring. Dropout rates were compared by demographic and clinical characteristics. Rate ratios (RR) and 95% confidence intervals (CI) were calculated for the association of factors with dropping out using a Poisson regression model. All factors for which univariate associations attained a significance of p < 0.1 on the log-likelihood test were included in an initial multivariable model. Factors were retained in the multivariable model if the p-value for inclusion using the log-likelihood test was ≤0.05. We calculated a Kaplan-Meier estimate of dropout rate over time for the participants in the cohort. We further performed a sensitivity analysis excluding seventy-two that never returned at all for follow up. Similar approaches as for the primary analysis were followed for the sensitivity analysis.

### Ethical considerations

The study was approved by the Research and Ethics Committee of the Uganda Virus Research Institute, and by the Uganda National Council of Science and Technology. Written informed consent was obtained from each participant before enrolment. Those who were confirmed to be HIV sero-positive were referred for HIV care and treatment at the appropriate facility.

## RESULTS

### Screening and enrolment

A total of 940 individuals were screened between January 2012 and February 2015; 279 were not enrolled (245 were at low risk for HIV, 8 were HIV-infected, and 26 were excluded for other reasons). We present data on 654 (69.6%) participants who were due for at least one follow-up visit.

### Baseline characteristics

The majority (61.4%) of the participants were male, Table 1. The mean age was 27.7 years (SD 6.9). Nearly half were married, most (71.3%) had attained primary education and engaged in fishing or related occupations (51.1%). The majority of the sample (76.9%) was of Christian faith and of Baganda ethnicity (44.0%). Only a third had lived in the fishing community for more than five years and about 50% had stayed away from home for at least two days in the last three months. Most (60.0%) reported having two to three sexual partners in the last three months and three quarters reported having a new sexual partner in the same period with 63.5% reporting condom use with this new partner. Majority of the participants 462(70.6) were recruited from Lambu fishing community.

**Table 1:** Baseline socio-demographic and behavioural characteristics of men and women enrolled in a longitudinal HIV vaccine preparedness study, Masaka, Uganda.

| Variable | All |  | Male |  | Female |  | P-value of gender difference |
| --- | --- | --- | --- | --- | --- | --- | --- |
|  | N=654 |  | N=402 |  | N=252 |  |  |
|  | n | (%) | n | (%) | n | (%) |  |
| Age (years) |  |  |  |  |  |  | 0.003 |
| 35+ | 248 | 38.0 | 132 | 32.8 | 116 | 46.0 |  |
| 25-34 | 282 | 43.0 | 186 | 46.3 | 96 | 38.1 |  |
| 18-24 | 124 | 19.0 | 84 | 20.9 | 40 | 15.9 |  |
| Current marital status |  |  |  |  |  |  | <0.001 |
| Single | 186 | 28.4 | 132 | 32.8 | 54 | 21.4 |  |
| Married | 319 | 48.8 | 217 | 54.0 | 102 | 40.5 |  |
| Separated/Widowed/divorced | 149 | 22.8 | 53 | 13.2 | 96 | 38.1 |  |
| Education level |  |  |  |  |  |  | 0.033 |
| More than primary | 125 | 19.1 | 66 | 16.4 | 59 | 23.4 |  |
| Primary | 466 | 71.3 | 301 | 74.9 | 165 | 65.5 |  |
| None | 63 | 9.6 | 35 | 8.7 | 28 | 11.1 |  |
| Occupation |  |  |  |  |  |  | <0.001 |
| Small scale business | 49 | 7.5 | 32 | 8.0 | 17 | 6.7 |  |
| Fishing/fishing related<sup>1</sup> | 334 | 51.1 | 280 | 69.6 | 54 | 21.4 |  |
| Services(Bar/lodge/restaurant/saloon) | 67 | 10.2 | 21 | 5.2 | 46 | 18.3 |  |
| Other(Peasant farmer/House wife) | 204 | 31.2 | 69 | 17.2 | 135 | 53.6 |  |
| **Religion** |  |  |  |  |  |  | 0.729 |
| Muslim | 151 | 23.1 | 91 | 22.6 | 60 | 23.8 |  |
| Christian | 503 | 76.9 | 311 | 77.4 | 192 | 76.2 |  |
| **Tribe/Ethnic group** |  |  |  |  |  |  | <0.001 |
| Baganda | 288 | 44.0 | 178 | 44.3 | 110 | 43.7 |  |
| Banyankole | 93 | 14.2 | 58 | 14.4 | 35 | 13.9 |  |
| Banyarwanda | 136 | 20.8 | 64 | 17.9 | 72 | 28.6 |  |
| Other | 137 | 21.0 | 102 | 25.4 | 35 | 13.9 |  |
| **Duration of stay in community (years)** |  |  |  |  |  |  | <0.001 |
| 0-< 1 | 164 | 25.1 | 57 | 14.2 | 107 | 42.5 |  |
| >1- 5 | 289 | 44.2 | 201 | 50.0 | 88 | 34.9 |  |
| > 5 | 201 | 30.7 | 144 | 35.8 | 57 | 22.6 |  |
| **Fishing Community** |  |  |  |  |  |  |  |
| Lambu | 462 | 70.6 | 279 | 69.4 | 183 | 72.6 | 0.776 |
| Kachanga | 68 | 10.4 | 45 | 11.9 | 23 | 9.13 |  |
| Kaziru | 32 | 4.9 | 21 | 5.2 | 11 | 4.3 |  |
| Other | 92 | 14.1 | 57 | 14.2 | 35 | 13.9 |  |
| **Away from home ≥ 2 nights, last 3 mos** | 299 | 45.8 | 221 | 55.1 | 78 | 31.0 | <0.001 |
| **Number of sexual partners, last 3 mos** |  |  |  |  |  |  | <0.001 |
| 1 | 215 | 32.8 | 80 | 19.9 | 135 | 53.6 |  |
| 2 to 3 | 333 | 60.0 | 230 | 57.2 | 103 | 40.9 |  |
| 4 or more | 106 | 16.2 | 92 | 22.9 | 14 | 5.6 |  |
| **New sexual partner, last 3 mos** | 480 | 74.1 | 324 | 81.2 | 156 | 62.7 | <0.001 |
| **Condom use with new partner, last 3 mos** | 305 | 63.5 | 205 | 63.3 | 100 | 64.1 | 0.859 |
| **Drank alcohol , in last month** |  |  |  |  |  |  |  |
| Never | 239 | 36.6 | 128 | 31.9 | 112 | 44.1 |  |
| Sometimes | 354 | 54.1 | 229 | 57.0 | 126 | 49.6 |  |
| Daily | 61 | 9.3 | 45 | 11.1 | 16 | 6.3 |  |
| **Drank alcohol before sex, in last month** |  |  |  |  |  |  | 0.005 |
| Never | 352 | 53.8 | 197 | 49.0 | 155 | 61.5 |  |
| Sometimes | 204 | 31.2 | 135 | 33.6 | 69 | 27.4 |  |
| Always | 98 | 15 | 70 | 17.4 | 28 | 11.1 |  |
| **Drug use, in last month** |  |  |  |  |  |  | <0.001 |
| None | 570 | 87.2 | 334 | 83.1 | 236 | 93.7 |  |
| Marijuana | 35 | 5.3 | 25 | 6.2 | 10 | 4.0 |  |
| Khat | 41 | 6.3 | 39 | 9.7 | 2 | 0.8 |  |
| Other | 8 | 1.2 | 4 | 1.0 | 4 | 1.6 |  |
| **Genital sores, last 3 mos, self-report** | 272 | 41.6 | 145 | 36.1 | 127 | 50.4 | <0.001 |
| **Laboratory confirmed syphilis** | 23 | 3.5 | 16 | 4.0 | 7 | 2.8 | 0.417 |
| **Circumcised** | -- | -- | 164 | 42.0 | -- | -- |  |

Mos-Month

More women (46.0%) than men (32.8%) were more than 35 years of age, p=0.003. More men than women reported being single (32.8% vs. 21.4%) and being divorced or widowed (38.1% vs. 13.2%), p<0.001. More women had attained more than primary education (23.4% vs. 16.4%, p=0.033). While most (69.6%) of the men were involved in fishing or fishing related occupations, most women (53.6%) were engaged in other activities, p<0.001. At enrolment, a higher proportion of men reported having been living in the community for more than 5 years (35.8%) compared to women (22.6%), p<0.001. More than half of men (55.1%) reported spending more than two nights away from home compared to women (31.0%), p<0.001. In the previous three months, more women (53.6%) than men (19.9%) reported having had only one sexual partner, p<0.001. More males (82.1%) reported having new partners in the last three months compared to females (62.7%), p<0.001. More males (17.4%) than females (11.1%) reported always drinking before sex, p=0.005.

### Study drop out

One hundred ninety-seven (30.1%) of participants dropped out of the study, (Figure 1), including 72 who did not return for the first follow up visit at month three. The dropout rate averaged over the entire 24-month study period was 25.3 / 100 PYO (95% CI: 22.0-29.1), Table 2. Excluding the 72 volunteers who never returned, the dropout rate was 17.1/100 PYO (95% CI 14.4-20.4). The most common reasons for dropping out included: moving away from study area (112), being untraceable (26), withdrawal from the study (31), and death (3). Fifteen participants were determined to be ineligible due to lower risk at the 12-month visit, and did not participate further (i.e. they were not counted among drop outs). During the two years, 45 participants became HIV-infected and were also not counted among drop outs. The HIV incidence amongst study participants in this cohort of 6.04 per 100 person years at risk (95% confidence interval: 4.36 – 8.37) has been reported previously (15).

**Figure 1.**
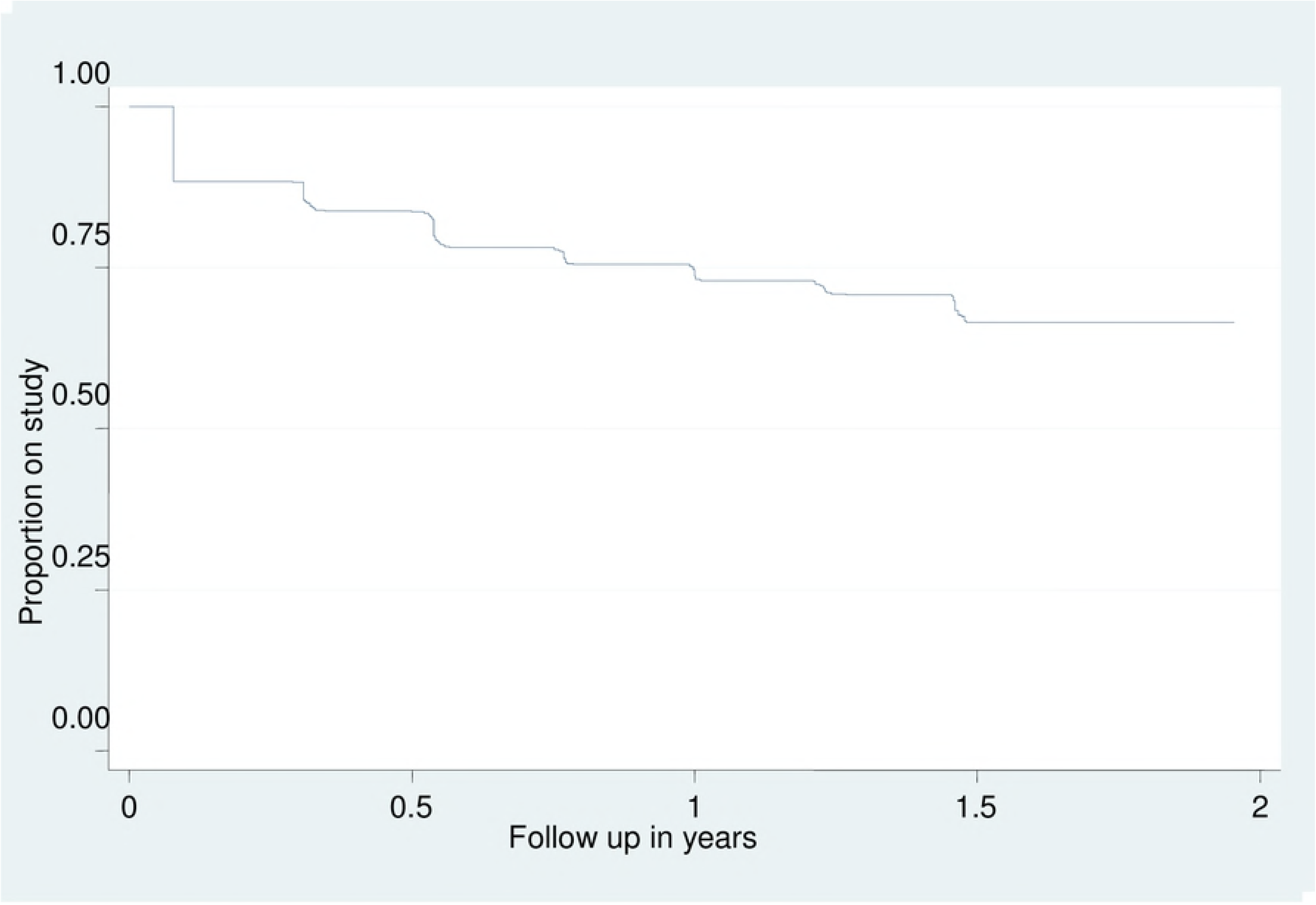
Kalpan Merier plot of volunteer attrition over time.

**Table 2:** Demographic and behavioural factors associated with study drop out (DO)

| Variable | N | DO | PYO | DO/100PO | RR(95% CI) | LRT P-value | aRR(95% CI) | LRT p-value |
| --- | --- | --- | --- | --- | --- | --- | --- | --- |
| **Overall 2 year drop out** | 654 | 197 | 778.9 | 25.3 |  |  |  |  |
| **Gender** |  |  |  |  |  |  |  |  |
| Male | 402 | 103 | 507.9 | 20.3 | Ref | <0.001 | Ref | 0.002 |
| Female | 252 | 94 | 270.9 | 34.7 | 1.7(1.29-2.26) |  | 1.56(1.12-2.18) |  |
| **Age (years)** |  |  |  |  |  |  |  |  |
| 35+ | 124 | 25 | 166.3 | 15 | Ref | 0.004 | Ref | 0.007 |
| 25-34 | 282 | 88 | 340.3 | 25.9 | 1.72(1.10-2.68) |  | 1.63(1.04-2.55) |  |
| 18-24 | 248 | 84 | 272.2 | 30.9 | 2.05(1.31-3.21) |  | 1.64(1.03-2.60) |  |
| **Current marital status** |  |  |  |  |  |  |  |  |
| Single | 186 | 64 | 217.4 | 29.4 | Ref | 0.107 |  |  |
| Married | 319 | 85 | 394.5 | 21.5 | 0.73(0.58-1.01) |  |  |  |
| Separated/Widowed | 149 | 48 | 166.9 | 28.8 | 0.98(0.67-1.42) |  |  |  |
| **Educational level** |  |  |  |  |  |  |  |  |
| More than primary | 125 | 39 | 156.0 | 25 | Ref | 0.013 | Ref | 0.01 |
| Primary | 466 | 131 | 563.3 | 23.3 | 0.93(0.65-1.33) |  | 1.06(0.74-1.52) |  |
| None | 63 | 27 | 59.5 | 45.4 | 1.82(1.11-2.97) |  | 2.02(1.23-3.31) |  |
| **Occupation** |  |  |  |  |  |  |  |  |
| Small scale business | 49 | 8 | 68.3 | 11.7 | Ref | 0.006 | Ref | 0.035 |
| Fishing/fishing related<sup>1</sup> | 334 | 99 | 413.2 | 24 | 2(0.99-4.21) |  | 1.98(0.95-4.11) |  |
| Services (Bar/lodge/restaurant/saloon) | 67 | 17 | 80.0 | 21.2 | 1.8(0.78-4.21) |  | 1.18(0.50-2.78) |  |
| Other | 204 | 73 | 217.3 | 33.6 | 2.9(1.38-5.95) |  | 2.05(0.98-4.31) |  |
| **Religion** |  |  |  |  |  |  |  |  |
| Muslim | 151 | 41 | 187.8 | 21.8 | Ref | 0.272 |  |  |
| Christian | 503 | 156 | 591.1 | 26.4 | 1.21(0.86-1.70) |  |  |  |
| **Tribe/Ethnic group** |  |  |  |  |  |  |  |  |
| Baganda | 288 | 83 | 339.2 | 24.5 | Ref | 0.762 |  |  |
| Banyankole | 93 | 29 | 103.1 | 28.1 | 1.15(0.75-1.75) |  |  |  |
| Banyarwanda | 136 | 39 | 170.3 | 23 | 0.94(0.64-1.37) |  |  |  |
| Other | 137 | 46 | 166.3 | 27.7 | 1.13(0.79-1.62) |  |  |  |
| **Duration of stay in community (years)** |  |  |  |  |  |  |  |  |
| >5 | 201 | 41 | 277.8 | 14.8 | Ref | <0.001 | Ref | 0.004 |
| 1 to 5 | 289 | 92 | 341.3 | 27 | 1.83(1.26-2.64) |  | 1.68(1.16-2.45) |  |
| 0-<1 | 164 | 64 | 160.0 | 40 | 2.71(1.83-4.02) |  | 2.22(1.46-3.38) |  |
| **Fishing community** |  |  |  |  |  |  |  |  |
| Lambu | 462 | 144 | 555.5 | 25.9 | Ref | 0.474 |  |  |
| Kachanga | 68 | 10 | 39.6 | 25.2 | 0.97(0.51-1.85) |  |  |  |
| Kaziru | 32 | 14 | 79.9 | 17.5 | 0.68(0.39-1.67) |  |  |  |
| Other | 92 | 29 | 103.8 | 27.9 | 1.07(0.72-1.61) |  |  |  |
| **Away from home ≥ 2 nights, last 3 mos** |  |  |  |  |  |  |  |  |
| No | 354 | 107 | 425.1 | 25.2 | Ref | 0.912 |  |  |
| Yes | 299 | 90 | 351.9 | 25.6 | 1.02(0.77-1.34) |  |  |  |
| **Number of sexual partners, last mos** |  |  |  |  |  |  |  |  |
| 0-1 | 215 | 64 | 261.7 | 24.5 | Ref | 0.565 |  |  |
| 2 to 3 | 333 | 97 | 396.8 | 24.4 | 0.99(0.73-1.37) |  |  |  |
| 4 + | 106 | 36 | 120.4 | 29.9 | 1.22(0.81-1.84) |  |  |  |
| **New sexual partner, last 3 mos** |  |  |  |  |  |  |  |  |
| No | 168 | 48 | 205.9 | 23.3 | Ref | 0.469 |  |  |
| Yes | 480 | 148 | 563.7 | 26.3 | 1.12(0.81-1.56) |  |  |  |
| **Condom with new partner, last 3 mos** |  |  |  |  |  |  |  |  |
| No | 175 | 60 | 198.7 | 30.2 | Ref | 0.322 |  |  |
| Yes | 305 | 88 | 365.0 | 24.1 | 0.79(0.57-1.11) |  |  |  |
| **Drank alcohol, last month** |  |  |  |  |  |  |  |  |
| Never | 293 | 80 | 280.5 | 22.9 | Ref | 0.41 |  |  |
| Sometimes | 354 | 98 | 418.9 | 19.2 | 0.82(0.61-1.10) |  |  |  |
| Daily | 61 | 19 | 79.4 | 15.3 | 0.84(0.51-1.38) |  |  |  |
| **Drank alcohol before sex, last mos** |  |  |  |  |  |  |  |  |
| Never | 239 | 113 | 407.8 | 27.7 | Ref | 0.368 |  |  |
| Sometimes | 354 | 58 | 254.3 | 22.8 | 0.82(0.59-1.13) |  |  |  |
| Always | 61 | 26 | 116.8 | 22.3 | 0.8(0.52-1.23) |  |  |  |
| **Illicit drugs use,in last month** |  |  |  |  |  |  |  |  |
| None | 570 | 169 | 680.5 | 24.8 | Ref | 0.754 |  |  |
| Khat | 35 | 11 | 46.8 | 23.5 | 0.94(0.51-1.74) |  |  |  |
| Marijuana | 41 | 14 | 42.8 | 32.7 | 1.32(0.76-2.27) |  |  |  |
| Other | 8 | 3 | 8.9 | 33.7 | 1.35(0.43-4.24) |  |  |  |
| **Genital sores, in last 3 months** |  |  |  |  |  |  |  |  |
| No | 382 | 114 | 454.7 | 25.1 | Ref | 0.884 |  |  |
| Yes | 272 | 83 | 324.1 | 25.6 | 1.02(0.77-1.36) |  |  |  |
| **Laboratory Confirmed Syphilis** |  |  |  |  |  |  |  |  |
| No | 591 | 180 | 695.1 | 25.9 | Ref |  |  |  |
| Yes | 63 | 17 | 83.7 | 20.3 | 0.78(0.48-1.36) |  |  |  |
| **Male circumcision** |  |  |  |  |  |  |  |  |
| Yes | 164 | 64 | 296.3 | 21.6 | Ref |  |  |  |
| No | 231 | 37 | 202.6 | 18.3 | 0.84(0.56-1.27) |  |  |  |

<sup>1</sup>Drying fish, salting or smoking fish, Mos-Month

Factors that remained independently associated with dropping out in the adjusted analysis included: gender, with females at increased risk of dropping out (adjusted rate ratio (aRR) =1.56; 95% CI 1.12-2.18) compared to males, education level, with participants having no education at all being twice more likely to drop out (aRR= 2.02; 95% CI: 1.23-3.31) than those with more than primary education (Table 2). Younger participants were at greater risk of dropping out, those aged 18-24 years old (aRR=1.64; 95% CI: 1.03-2.06) and 25-34 years old (aRR = 1.63; 95% CI: 1.04-2.55) compared to those aged 35+ years. Short duration of stay in community, participants who reported to have lived in the fishing community for one year or less were two times more likely to drop out (aRR=2.22; 95% CI: 1.46-3.38) compared to those who had spent over 5 years. Occupation was also independently associated with drop out with participants engaged in fishing (aRR=1.98; 95% CI: 0.95-4.11), services (aRR=1.18; 95% CI: 0.50-2.78) and other (aRR=2.05; 95% CI: 0.98-4.31) being more likely to drop out than those involved in small scale business.

Compared to those who completed at least one follow up visit, those who never returned for follow up were younger (mean age of 25.5 vs. 28.0; p=0.004) and more likely to have had no education (19.4% vs 8.4%, p=0.010) and to have lived for less than a year in the community (45.8% vs 22.5%, p<0.001).

## DISCUSSION

In this HIV vaccine preparedness open cohort study among fisher folk, we observed a dropout rate of 25.3/100 person years of observation over the 2-year duration of follow-up, or 17.1/100 PYO if you exclude those who never returned for any follow up visits. The findings from our study build on reports from previous work conducted on these and other similar populations around Lake Victoria. Retention rates of 85% (12) and 77% (11) were reported in two other fisher folk cohorts in which participants were followed up in community-based clinics. Our study differs from these studies in three main aspects firstly, these studies (11–13) differed from our study in that they reported their retention as proportions, while our study is reporting a dropout rate over the two year period. Though the difference may be modest, a proportion will typically overestimate drop out compared to a rate (i.e., 30.1% of our study participants dropped out, but our rate was 25.3 drop outs per 100 PYO). Secondly the study retention for some of the studies (11, 12) were performed in the fishing communities whereas our study activities were at a distant study clinic, and thirdly some also (11, 13) reported their retention over a period of 1 year compared to 2 years in our study.

The observed high rate of drop out at the month 3 may point to the fact that some participants were willing to enrol but were simply unwilling to return for any follow up. This finding suggests for the adoption of a lead-in type of enrolment scheme for clinical trials involving participants from these fishing communities.

Even though we provided reimbursement for transportation and time, as well as free STI treatment, this may not have been adequate to overcome the hurdles of having to travel to a distant research site. Distance from home could have affect retention in follow up. In a cohort study conducted among discordant couples in Kenya, it was discovered that participants who were living 5-10 kilometres away from the clinic were twice as likely to have follow-up interruptions compared to those living less than 5 kilometres(16). Although no differences were observed in dropout rates between the different fishing communities, it is possible that the distance from the fishing communities to the clinic could have contributed to the drop out.

We found that persons who had lived in the community for a relatively short time were more likely to drop out of the study. This findings is consistent with those of previous studies in Uganda fishing communities (11) (12). It is likely that being in a community for a shorter period of time is a proxy for people who are more likely to be migratory and who tend to move between different islands and landing sites usually following the seasonal fishing seasons (17). I think this would be a good spot to talk about enrolment village and distance to masaka study clinic, and if we did/did not see any relationships

Women may also be less likely to be retained, because they usually have time demanding roles within a family including care for sick family members, provision of care for children and food. It is possible too that some women are commercial sex workers and who might move from place to place depending on the demand for their services. Women will therefore find difficulty in taking time off to come to the clinic. Since it is important to ensure that men and women are equally engaged in HIV vaccine efficacy trials, efforts need to be made to increase female enrolment and retention in clinical trials within these communities. Participation of women from the fishing communities can potentially be improved by involvement of individuals or groups in the community who have ample rapport with them, such as, their spouses, brothel owners, bar and lodge owners.

The observation that younger participants are more likely to drop out. This finding is similar to what was observed in the previous studies (11, 12) and may be due to the fact that young people possibly haven’t settled in any stable relationship/occupation and are still moving in search of greener pastures.

The strength of this study included longer duration of follow up with frequent clinic visit in a set up with an already established infrastructure suitable for HIV vaccine efficacy trials. This builds on the previous studies that provided retention data but from follow up at clinics established within the fishing communities.

Our study had some limitations. Firstly, we were often not able to document the specific reasons why participants dropped out, other than some data on relocation. This information could have probably provided more insight on the specific reasons for dropping out. Secondly, at the initial contact with potential participants in the fishing communities, we did not collect data on those who attended meetings but who did not later attend screening visits at the Masaka study clinic, nor did we collect data on those who attended screening at the clinic but were not enrolled (aside from reason they were not enrolled), thus we were unable to make comparisons between enrolled and non-enrolled persons. Thirdly, we did not collect data on breastfeeding or pregnancy among the females, which may have also been barriers for participation and retention, since these may affect ones travel to and from the study clinic.

## Conclusions

Although we demonstrate ability to enrol participants in this longitudinal cohort, the high dropout rate especially in the first three months is a point of concern. Inclusion of volunteers into clinical trials to evaluate products such as vaccine candidates would require delayed enrolment and consider an initial pre-screening period of approximately 2-3 months, requiring potential volunteers to attend more than one screening visit to establish commitment. Particular attention for retention should be paid to the participant categories identified as being a risk of dropping out such as female, younger, lower educated and participants with short duration of stay in the community.

## Acknowledgement

This work was funded by IAVI with the generous support of USAID and other donors; a full list of IAVI donors is available at www.iavi.org. The contents of this manuscript are the responsibility of IAVI and co-authors and do not necessarily reflect the views of USAID or the US Government. We also wish to acknowledge support from the University of California, San Francisco’s International Traineeships in AIDS Prevention Studies (ITAPS), U.S. NIMH, R25MH064712. As part of the ITAPS we thank the helpful reviews of Professor Rhoderick Machekano, and peer reviews of Drs Huub Gulderblom, Carol Camlin and Mi Suk. We recognise the logistical support of the Administration team at UCSF-ITAPS Dr Jeffery Mendel, Dr Debbie Brickley and Ritu Sehgal.

## Notes

**Conflicts of interest**: No conflict of interest

